# Mitochondrial ascorbate synthesis acts as a pro-oxidant pathway and down-regulate energy supply in plants

**DOI:** 10.1101/825208

**Authors:** Luis Miguel Mazorra Morales, Gláucia Michelle Cosme Silva, Diederson Bortolini Santana, Saulo F. Pireda, Antonio Jesus Dorighetto Cogo, Angelo Schuabb Heringer, Tadeu dos Reis de Oliveira, Ricardo S. Reis, Luís Alfredo dos Santos Prado, André Vicente de Oliveira, Vanildo Silveira, Maura Da Cunha, Claudia F. Barros, Arnoldo R. Facanha, Pierre Baldet, Carlos Bartoli, Marcelo Gomes da Silva, Jurandi Gonçalves de Oliveira

## Abstract

Attempts to improve the ascorbate (AsA) content of plants are still dealing with the limited understanding of why exists a wide variability of this powerful anti-oxidant molecule in different plant sources, species and environmental situations. In plant mitochondria, the last step of AsA synthesis is catalyzed by the enzyme L-galactone-1,4-lactone dehydrogenase (L-GalLDH). By using GalLDH-RNAi silencing plant lines, biochemical and proteomic approaches, we here discovered that, in addition to accumulate this antioxidant, mitochondria synthesize AsA to down-regulate the respiratory activity and the cellular energy provision. The work reveals that the AsA synthesis pathway within mitochondria is a branched electron transfer process that channels electrons towards the alternative oxidase, interfering with conventional electron transport. It was unexpectedly found that significant hydrogen peroxide is generated during AsA synthesis, which affects the AsA level. The induced AsA synthesis shows proteomic alterations of mitochondrial and extra-mitochondrial proteins related to oxidative and energetic metabolism. The most identified proteins were known components of plant responses to high light acclimation, programmed cell death, oxidative stress, senescence, cell expansion, iron and phosphorus starvation, different abiotic stress/pathogen attack responses and others. We propose that changing the electron flux associated with AsA synthesis might be part of a new mechanism by which the L-GalLDH enzyme would adapt plant mitochondria to fluctuating energy demands and redox status occurring under different physiological contexts.

## INTRODUCTION

In plants, the mitochondrial electron transport chain (mETC) consists of a series of electron transporters that function to oxidize reducing equivalents, NADP(H) and FADH2 (Schertl and Braun, 2014). A widely accepted model about the electron transfer is that the electrons normally enter via complex I (NADH:ubiquinone oxidoreductase) or through a diversity of “alternative” NAD(P)H dehydrogenases using flavin mononucleotide (FMN) as electron acceptor (Pineau et al., 2005).

Alternatively, complex II (succinate:ubiquinone oxidoreductase) and other dehydrogenases such as glyceraldehyde 3-phosphate dehydrogenase (G3-PDH), the “electron transfer flavoprotein-ubiquinone oxidoreductase” (ETFQ-OR) and the proline dehydrogenase (ProDH) supply electrons to mETC but via flavin dinucleotide (FAD) (Sweetlove et al., 2010). These oxidation reactions are all coupled to reduction of the ubiquinone (UQ) to ubiquinol (UQH_2_) (Schertl and Braun, 2014).

To accomplish ubiquinol re-oxidation, two routes have been proposed. The cytochrome c oxidase pathway (COX), in which electrons in UQH_2_ are then passed to complex III (ubiquinone:cytochrome c oxidoreductase), which reduces cytochrome c (Cytc) and oxidizes UQH_2_ and subsequently the reduced Cytc is re-oxidized by complex IV (cytochrome c oxidase) with dependence of electron reduction of O_2_ to H_2_O (Millar et al., 2011). The other, the so-called “alternative” oxidase pathway (AOX), directly oxidizes UQH_2_ coupled with the reduction of O_2_ to H_2_O (Vanlerberghe, 2013). Thus, the AOX introduces a branch in the mETC and consequently a regulatory point that allows the partition of electrons between both pathways.

The electrons channeled via respiratory complexes (I, III and IV) are coupled to the pumping of H^+^ through the inner-mitochondrial membrane (Millar et al., 2011). Pumping of H^+^ results in a transmembrane proton gradient, which is required for generation of adenosine triphosphate (ATP) from ADP and Pi via complex V (H^+^-ATP synthase), a process termed as oxidative phosphorylation.

The plant mitochondrial electron transport greatly depends upon AOX pathway. Indeed, the AOX protein is the most highly regulated component of mETC at transcriptional and translational level (Vanlerberghe et al., 2016; Dahal and Vanlerberghe, 2017). When ATP demand is low, ATP synthesis needs to be decreased and the electrons would flow via AOX pathway, it bypasses the H^+^-pumping protein complexes III and IV, reducing the potential of mitochondrial ATP production (Wagner & Wagner, 1995; Millar et al., 2011; Vanlerberghe, 2013). For example, high AOX pathway in light would act in coordination with photosynthesis (an ATP source) to optimize energy metabolism (Nunes-Nesi et al., 2011).

Other role for AOX pathway is to avoid the over-flux through COX pathway and consequently ROS over-production (Vanlerberghe et al., 2016; Dahal and Vanlerberghe, 2017). When ATP supply through COX pathway is uncoupled from energy demand, it leads to the over-reduction of mETC and the formation of partially reduced forms of electron carriers such as flavins (FAD, FMN) and ubiquinone, which can potentially react with oxygen (O_2_) to form reactive oxygen species (ROS), e.g. O_2_^−^, H_2_O_2_, ^1^O_2_ (Mailloux and Harper, 2011). If ROS are not effectively scavenged, they would generate oxidative damage (Noctor and Foyer, 2016). ROS can also have positive signaling roles at low concentrations and therefore control of ROS load by AOX has been involved in ROS signaling, programmed cell death, abiotic/biotic stress tolerance and plant growth (Vanlerberghe, 2013).

Ascorbate is an abundant molecule in plants, which plays multiple roles as antioxidant, pro-oxidant and co-factor for multiple enzymes in plants and mammals reviewed in Smirnoff, 2018. AsA is ultimately synthesized by plant mitochondria through another dehydrogenase, the L-galactone-1,4-lactone dehydrogenase (L-GalLDH EC 1.3.2.3). This enzyme is a FAD-dependent dehydrogenase that catalyzes the L-Galactone-1,4-lactone (L-GalL) oxidation. The plant L-GalLDH enters electrons directly to mETC through cytochrome c (Bartoli et al., 2000). L-GalLDH expression and AsA synthesis are under diurnal control (Tamaoki et al., 2003) and are up-regulated by light in an AOX-dependent manner (Bartoli et al., 2006). Conversely, L-GalDH expression and AsA synthesis are down-regulated in dark and with ageing (Tamaoki et al., 2003). Beyond its roles in AsA synthesis and in the assembly of complex I (Schertl et al., 2012), the significance of L-GalLDH as component of mETC is still unclear.

The analysis of several AsA-deficient mutants with defects in key points of AsA biosynthesis pathway highlights different roles of AsA through plant lifecycle. AsA-biosynthesis mutants have showed alterations in flowering and senescence (Kotchoni et al., 2009), enhanced sensitivity to high light, salt, UV-B radiation and extreme temperatures (Smirnoff, 2011), biotrophic pathogen resistance (Mukherjee et al., 2010), tolerance to postsubmergence reoxygenation (Yuan et al., 2017); hormonal control (Foyer et al., 2007), sucrose and iron uptake (Grillet et al., 2014) and plant growth (Alhagdow et al., 2007). Interestingly, several physiological effects seemed to be independent of AsA deficiency (Smirnoff, 2018); however, the causes of this independence are unclear. The analysis of AsA synthesis-altered plants, specifically in the mitochondrial L-GalLDH expression, would offer a valuable tool to explore the roles of the last step of AsA synthesis into mitochondria and the impact on plant growth, development and stress tolerance.

We here revealed that mitochondrial AsA synthesis down-regulates energy supply and generates hydrogen peroxide accumulation. This pathway is a branched electron transfer process that channels electrons towards alternative oxidase. The proteomic of tissues with enhanced AsA synthesis reveals proteins related to oxidative and energetic homeostasis, suggesting that mitochondrial AsA synthesis leads to a global switch in redox and energy loads. These findings establish a new pro-oxidant function for AsA synthesis beyond producing this powerful anti-oxidant.

## RESULTS

### The alternative respiration is modulated by mitochondrial ascorbate synthesis

To answer the question how the L-GalLDH enzyme affects the mitochondrial electron transport chain (mETC), we examined the effects of respiratory inhibitors on mitochondrial respiration of RNAi-plant lines harboring silenced L-GalLDH activity. As expected, the mixture of AOX and COX inhibitors (5 mM SHAM and 3 mM NaN3) decreased the oxygen uptake rate of leaf mitochondria purified from wild type plants and L-GalLDH-RNAi plant lines (Figure 1A). Residual oxygen uptakes were observed in presence of both inhibitors. When leaf mitochondria were pre-treated with the L-GalLDH substrate (5 mM L-GalL), absolute respiration was greatly reduced and was not sensitive to the mixture of both inhibitors in wild type mitochondria. Nonetheless, a significant blockage of respiration occurred in the L-GalL-treated leaf mitochondria from L-GalLDH-RNAi plant lines (Figure 1A), which suggest that part of the AOX and COX pathways is active. The western blot analysis showed lower levels of L-GalLDH protein abundance in both L-GalLDH-RNAi plant lines (~21% and ~63% of Wt for 8-14 and 5-13 plant lines, respectively), which resulted in decreased L-GalLDH activity (Figure 1B). Notably, the level of L-GalLDH suppression but not the enzyme activity was more marked in the L-GalLDH-RNAi line 8-14 as compared to 5-13 line.

**Figure 1A.**
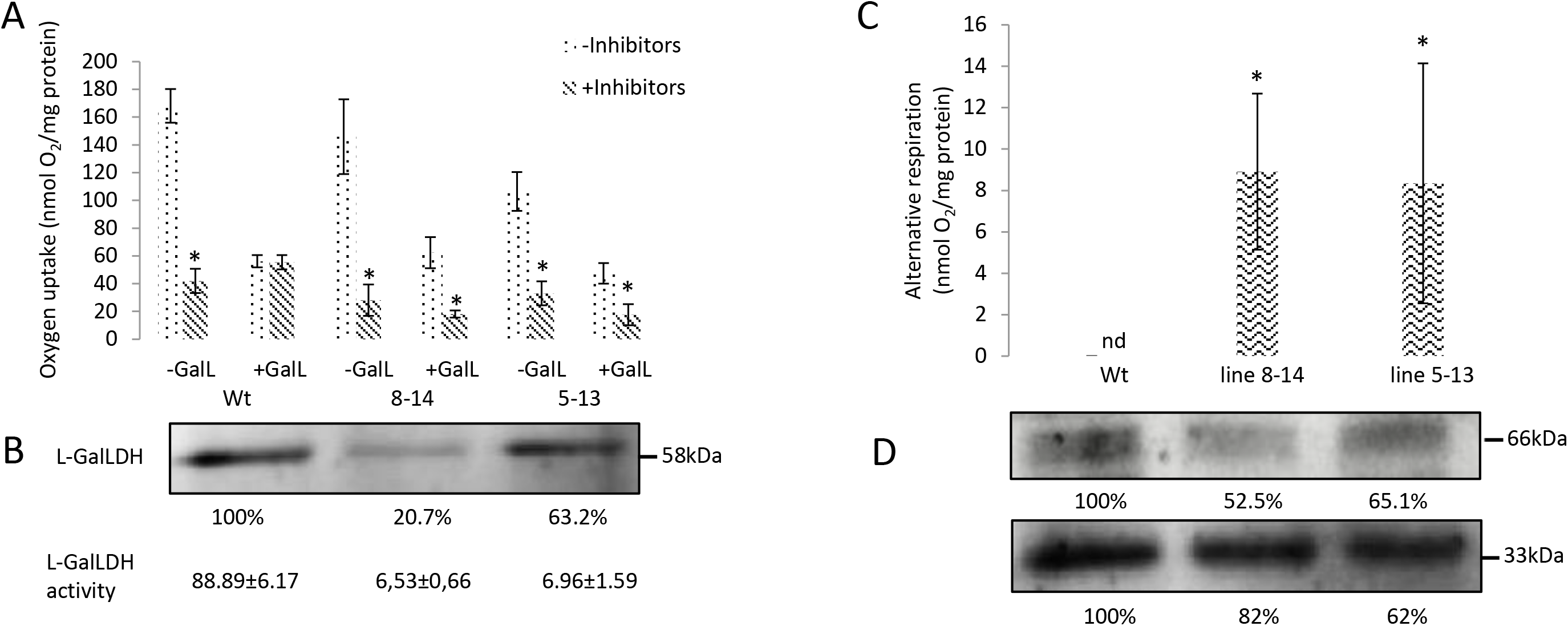
NADH-driven respiration of leaf mitochondria purified from 30-days-old wild type, 8-14 and 5-13 transgenic lines. Mitochondrial preparations were pre-treated or not with 5 mM L-GalL and the oxygen uptake rates were determined following the addition of 10 mM malate. Then, the respiration was blocked by a mixture of respiratory inhibitors 5 mM SHAM and 3 mM NaN3. **1B**. Immunoblot of L-GalLDH in leaf mitochondria from 30-days-old wild type, 8-14 and 5-13 transgenic lines detected by Western Blot using anti-L-GalLDH (Agrisera). Relative abundance, expressed as % of wild type signal, was obtained by densitometry. L-GalLDH activity (measured as rate of Cytc reduction) was determined in the mitochondria from 30-days-old wild type, 8-14 and 5-13 transgenic lines. Values represent means ± standard error and asterisks represent significant differences between inhibited and non-inhibited reactions analyzed by one-way ANOVA following by Tukey test (P < 0.05). Measurements from three independent mitochondrial preparations (n=3). **1C**. Rates of oxygen uptake SHAM-sensitive and NaN3-resistant (alternative respiration) determined in mitochondria from wild type, 8-14 and 5-13 transgenic lines. Asterisks represent significant differences of alternative respiration of transgenic leaf mitochondria when compared to wild type by one-way ANOVA following by Tukey test (P < 0.05). **1D**. Immunoblot of AOX detected when mitochondrial proteins are loaded without the reducing agent of free sulphydryl residues (2-mercaptoethanol) into sample buffer. A 66 kDa protein was detected by Western Blot using anti-AOX antibodies (Agrisera) in leaf mitochondria purified from 30-days-old wild type, 8-14 and 5-13 plants. Immunoblot of AOX (molecular weight of about 33 kDa) obtained when the reducing agent was added into sample buffer and subsequent detection with anti-AOX antibodies. Equal loading of gels was checked by Ponceau staining. Relative abundances were assessed by quantification of signals through densitometry and expressed as % of wild type level.

To further explore the causes of the differences in respiratory rates, we analyzed the flow of electrons through the AOX pathway in the presence of L-GalLDH substrate. Clearly, the Figure 1C shows that leaf mitochondria of the L-GalLDH-RNAi plants had significant alternative respiration ~9 nmol O_2_/mg protein was resistant to NaN3 and sensitive to SHAM in presence of L-GalL. However, it was not detected in wild type mitochondria (Figure 1C). The SDS-PAGE electrophoresis of mitochondrial proteins and subsequent inmunobloting with anti-AOX antibodies allowed the detection of a band of ~66 kDa in addition to trace level of ~33 kDa protein. However, when the samples for SDS-PAGE electrophoresis were prepared with the reducing agent, 2-mercaptoethanol, into the sample buffer, the band of ~33 kDa intensified and that of ~66 kDa virtually disappeared (Data not shown). Because the 2-mercaptoethanol reduces disulfide bond linkages within proteins, the presence of 33 kDa AOX protein could result from the break of the intermolecular disulfide bond in the 66 kDa AOX dimer by this reducing agent. Consistently, previous works showed the existence of a single 33 kDa AOX isoform in tomato mitochondria (Holtzapffel et al., 2002) and the dimerization of the 33 kDa AOX isoform by the oxidant, diamide (Holtzapffel et al., 2002). Most importantly was that the quantification by densitometry of independent gels (with equal loading of proteins, supplementary data I) showed higher amount of AOX (for both the 66 kDa and 33 kDa AOX) in wild type plants while it was lower in L-GalLDH-RNAi plant lines (Figure 1D). Taken together, these data suggest that the AOX molecules in wild type mitochondria are significantly inhibited by L-GalL due to the L-GalLDH activity. In the plant lines, the suppression of L-GalLDH enzyme could, in turn, prevent AOX inhibition by L-GalL. Interestingly, the 8-14 plant line, which has the higher L-GalLDH suppression (Figure 1B) showed the lower level of oxidized AOX (Figure 1D).

We examined the respiratory capacity of mitochondria purified from heterotrophic tissues (fruits) of other plant species and using other respiratory substrates and inhibitors. The mitochondrial preparations of fruit purified with a Percoll density-gradient method had intact mitochondria (≥80% integrity). The content of mitochondria (based on mitochondrial protein) ranged from 1 to 3.4 mg protein. The oxygen uptake of papaya, strawberry and tomato fruit mitochondria was blocked by respiratory inhibitors (supplementary data IIA). However, respiration was insensitive to inhibitors in the presence of L-GalL (supplementary data IIA). Moreover, when energizing mitochondria with other substrates that enter electrons through complexes I (malate, glutamate) or II (succinate), the alternative respiration was blocked by L-GalL (supplementary data IIB). Clearly, the insensitivity of oxygen uptake to inhibitors (supplementary data IIA) and the loss of alternative respiration in presence of L-GalL (supplementary data IIB) were responses in fruit mitochondria that resembled to those found in wild type tomato leaf mitochondria treated with L-GalL (Figure 1A and 1C). It supports the hypothesis of that the L-GalLDH activity down-regulates mitochondrial electron flux by inhibiting the alternative oxidase pathway. Clearly, this is a general effect in both autotrophic and heterotrophic plant tissues.

To get further insights about the mechanism inactivating AOX pathway, we adopt papaya fruit mitochondria as model because their ability to synthesize AsA and the significant bulk of active mitochondria with high AOX capacity that can be easily obtained our results here and (Oliveira et al., 2015). Respiration and ascorbate production of papaya mitochondria were stimulated by increasing L-GalL concentrations up to about 5mM. Higher concentrations were progressively inhibitory, being the respiratory activity more sensitive to the inhibition by the substrate concentration (supplementary data III).

### The alternative oxidase but not the Cytc oxidase is critical for AsA biosynthesis

By using inhibitors that target specific points in the mETC, we analyzed the possible role of terminal oxidases during mitochondrial AsA synthesis. The current Bartoli’s model explaining mitochondrial AsA synthesis implies that Cytc and Cytc oxidase are absolute requirements for AsA production (Bartoli et al., 2000). As Cytc oxidase re-oxidizes Cytc quickly, the L-GalLDH activity, which was measured as rate of Cytc reduction, is assayed in presence of Cytc oxidase inhibitor. It was confirmed that the treatment of mitochondria with the inhibitor of Cytc oxidase (NaN3, azyde) led to over-accumulation of reduced Cytc (~6 μmol cytc.min^−1^mg protein^−1^, Figure IIB), consistent with a lower Cytc re-oxidation by this terminal oxidase. However, azyde-treated mitochondria still maintained a little capacity to synthesize ascorbate (~0.35 μg AsA mg protein^−1^, Figure IIA). This suggested that part of AsA synthesis could be independent of Cytc oxidase. On the other hand, the addition of the inhibitor of AOX pathway (SHAM) affected drastically the Cytc reduction by L-GalL (<1 μmol cytc.min^−1^mg protein^−1^, Figure IIB), and provoked a very low level of AsA content (Figure IIA), it suggests that SHAM limits electron flux through Cytc.

**Figure 2A.**
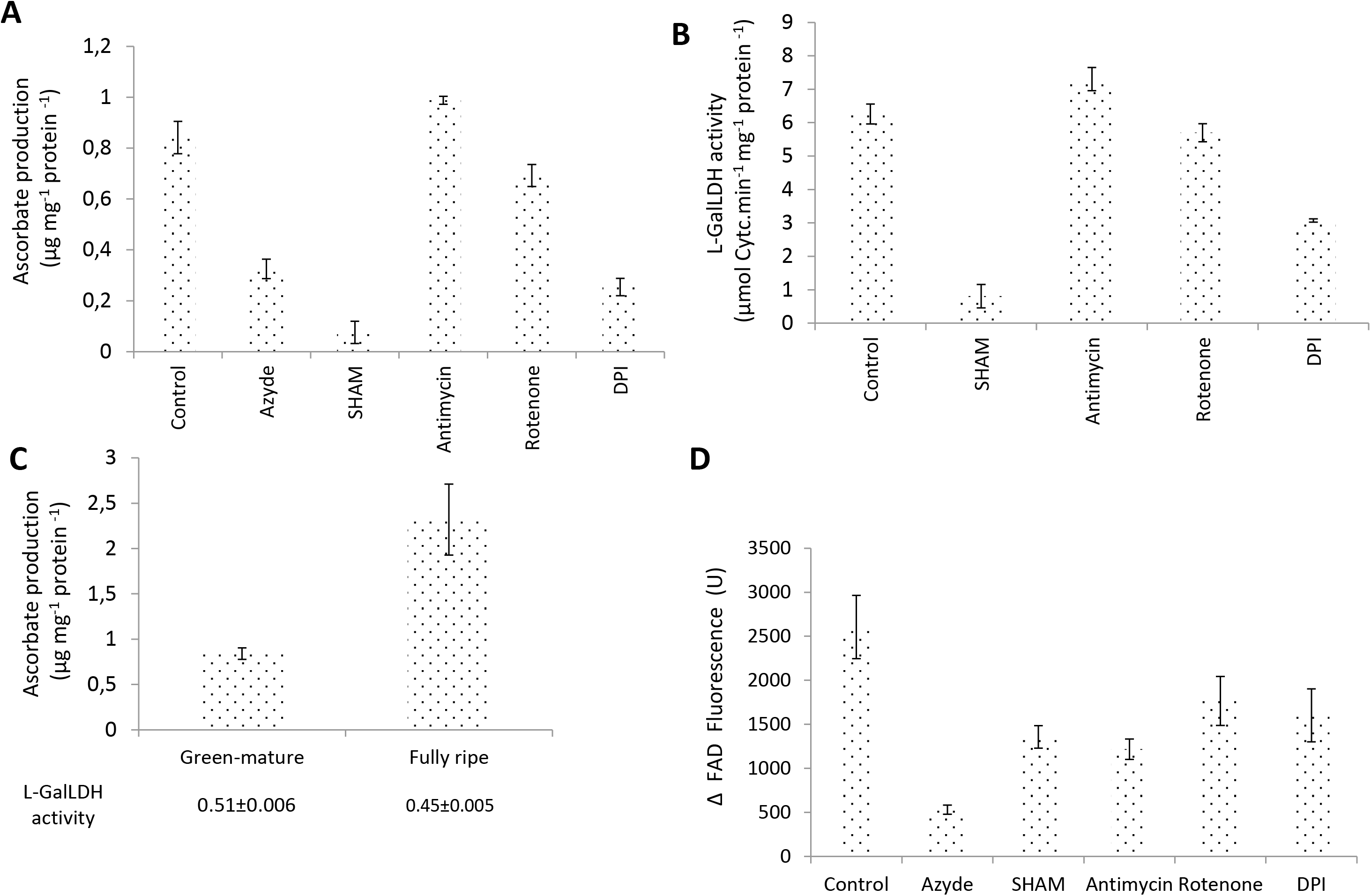
Ascorbate production capacity measured in pure mitochondria treated or not with inhibitors (3 mM azyde, NaN3, 1mM SHAM, 2 μM antimycin A, 20 μM rotenone, and 5 mM DPI). Ascorbate synthesis was initiated in presence of 5 mM L-GalL. **2B**. Activity of L-GalLDH enzyme (assessed as capacity to reduce Cytc) determined in purified mitochondria incubated with 1mM SHAM, 2 μM antimycin A, 20 μM rotenone, and 5 mM DPI). Cytc reduction was started by adding 5mM L-GalL in presence of the Cytc oxidase inhibitor, azyde. **2C**. Ascorbate production and L-GalLDH activity determined in mitochondria from green-mature and full-ripe papaya fruit. **2D**. Ascorbate synthesis-dependent changes in flavin adenine dinucleotide (FAD) fluorescence measured in pure mitochondria incubated with the inhibitor compounds indicated above in figure 2A. AsA synthesis was started with 5mM L-GalL. Bars represent means ± standard error from at least three independent experiments.

As AOX gene expression and capacity increase during papaya fruit ripening (Oliveira et al., 2015),we comparatively analyzed the L-GalLDH activity between green-mature and fully ripe papaya fruit. The mitochondria from ripe fruit showed lower L-GalLDH activity but had increased AsA synthesis capacity (Figure IIC).

In addition, other inhibitors also showed significant effects during AsA synthesis. Mitochondria treated with antimycin A, an inhibitor of complex III, showed the higher value of Cytc reduction (~7 μmol cytc.min^−1^mg protein^−1^) (Figure IIB) and favored AsA synthesis (Figure IIA). Moreover, when complex I was inhibited with rotenone, Cytc reduction and AsA synthesis were still maintained at levels similar to control (Figures IIB and IIA). By contrast, the DPI, an inhibitor of flavin-oxidases, affected markedly Cytc reduction and AsA synthesis (Figures IIB and IIA).

### Cytc oxidase is a main factor affecting FAD recycling during AsA synthesis

We followed changes in fluorescence of exogenously supplemented FAD in the presence of L-GalL and then assessed the effect of inhibitors on such changes. It was recorded a variation of FAD fluorescence (about 2500 Units) following incubation with L-GalL (Figure 2D). It indicates that FAD redox state changes during AsA synthesis. All inhibitors tested in this study decreased the effect of L-GalL in FAD fluorescence, having the Cytc oxidase inhibitor, azyde, the highest effect (Figure 2D). These data may suggest that the FAD redox state during AsA synthesis is basically controlled by Cytc oxidase, but other respiratory components could be also involved, albeit indirectly.

### Mitochondrial uncoupling and ROS over-production are associated with low AOX capacity during AsA synthesis

Given that alterations of mETC and AOX pathway may affect the pumping of H^+^ and the mitochondrial coupling (Millar et al., 2011), we explored if mitochondrial oxidative phosphorylation is also affected during AsA synthesis. As expected, there was a 19% of membrane depolarization (based on the respiratory increase induced by the uncoupling agent, CCCP) in NADH-respiring mitochondria. However, mitochondrial respiration was insensitive to CCCP in presence of L-GalL (Table I), suggesting that the generation of the proton gradient is affected. Moreover, both the phosphorylation efficiency, measured as ADP:O ratio and the mitochondrial coupling efficiency, determined as RCR, decreased in presence of L-GalL (Table I). Intriguingly, these L-GalL-dependent alterations correlated with higher H^+^-ATPase activity of complex V and an unexpected higher mitochondrial capacity to reduce NAD^+^ into NADH (Table I). As the ubiquinone redox state and the mitochondrial energy production are regulated by AOX pathway (Vanlerberghe, 2013), we analyzed possible changes in ubiquinone redox state and the mitochondrial ability to synthesize ATP. It was noted that, in the presence of L-GalL, the mitochondrial capacity to maintain UQ in its reduced state (UQH2) enhanced (about three times more reduced ubiquinone in the L-GalL treatment than in control). Besides, the mitochondrial ATP synthesis capacity was inhibited by L-GalL (Table I). These results suggest that AsA synthesis could cause the over-reduction of mETC and UQ pool, resulting in a decrease (~20%) in ATP synthesis capacity.

**Table I.** Effect of L-GalL on the mitochondrial ability to perform electron transport and coupled ATP synthesis.

|  | Treatments |  |
| --- | --- | --- |
|  | Control | + 5mM Gall |
| **Phosphorylation efficiency<sup>\*</sup>** | 3.06±0.15 | 0.63±0.16 |
| **Coupling efficiency** | 2.93±0.17 | 1.61±0.14 |
| **Depolarization** | 119% | 0% |
| **NADH production** | 0.021±0.0081 | 0.240±0.1731 |
| **H<sup>+</sup>-ATPase activity** | 208.07±29.87 | 365.46±78.74 |
| **UQH<sub>2</sub>/UQ ratio** | 2.49±0.81 | 7.05±0.24 |
| **ATP synthesis capacity** | 0.0373±0.0012 | 0.0298±0.0021 |

<sup>\*</sup>Efficiency of phosphorylation and coupling were determined recording the O<sub>2</sub> uptake rate of the coupled mitochondria in non-phosphorylating and phosphorylating states and expressed as ADP:O ratio and the respiratory control rate (RCR), respectively. Membrane depolarization was assessed by measuring the percentage of increase in the oxygen uptake induced by a respiratory uncoupler. Mitochondrial potential to reverse the electron flux was determined by quantifying the capacities to convert exogenous NAD<sup>+</sup> into NADH and to hydrolyze ATP in ADP and Pi. The capacity to alter the ubiquinone redox state was measured by the conversion of exogenously added oxidized ubiquinone into its reduced form and expressed as the ubiquinol/ubiquinone ratio. ATP synthesis capacity was assessed by quantifying the increase in the synthesis of ATP after adding ADP. All values are means± standard error from three independent mitochondrial preparations.

As AOX is an important ROS scavenger, we hypothesized that the inactivation of AOX pathway during AsA synthesis would enhance ROS. By measuring the H_2_O_2_ level, using the Amplex Red method, an increased H_2_O_2_ formation was detected within 5-15 min following incubation of mitochondria with L-GalL, having maximal H_2_O_2_ increases between 5-20 mM L-GalL whereas response was extremely low or non-detected at concentrations below 5 mM L-GalL (supplementary data IV).

We explored the possible sources of mitochondrial ROS during AsA synthesis. Figure IIIA shows the increase in H_2_O_2_ fluorescence (~50% above control) in mitochondria treated with 5mM L-GalL. This H_2_O_2_ fluorescence was maintained by AOX inhibitor (SHAM) whereas it was slightly higher than control in presence of Cytc oxidase inhibitor (azyde). Unexpectedly, relative H_2_O_2_ fluorescence was not further increased with the respiratory inhibitors rotenone (complex I), antimycin A (complex III), and DPI (flavin-oxidase inhibitor) (Figure IIIA).

**Figure 3A.**
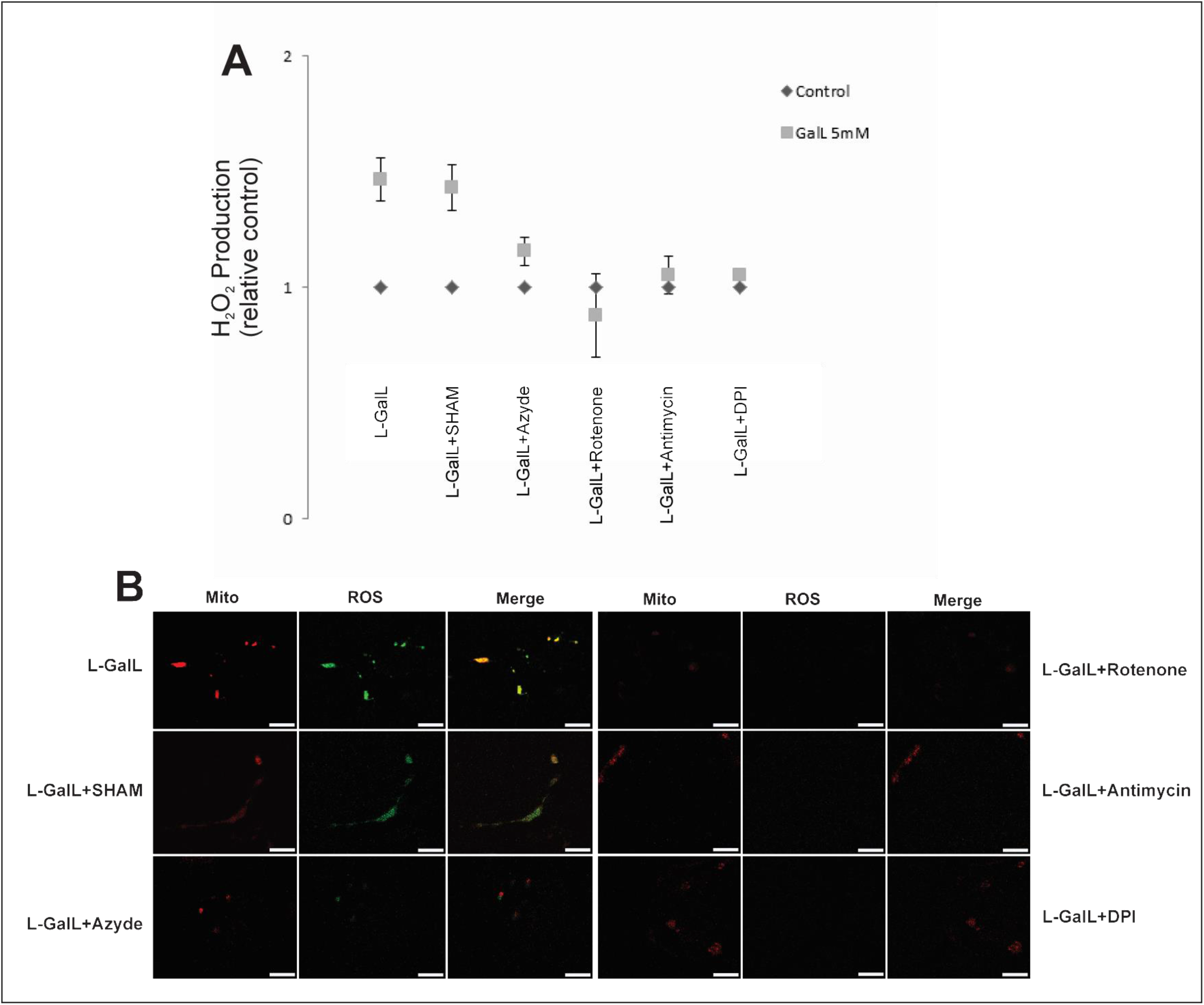
Induction of hydrogen peroxide (H_2_O_2_) production by 5 mM L-GalL in pure mitochondria treated with 1 mM SHAM, 3 mM azyde, 20 μM rotenone, 2 μM antimycin A, and 5 mM DPI. H_2_O_2_ content was quantified using Amplex Red/horseradish peroxidase (HRP) assay and relative H_2_O_2_ level for all treatments was normalized to their corresponding controls without L-GalL. Bars are means ± standard error (n=3). **3B**. Representative confocal images of co-localization of ROS and mitochondria stains in fruit mesocarp tissue incubated with 5 mM L-GalL and the same inhibitor compounds as above. Mitochondria were localized with Mito-Tracker Red and ROS detection was performed with (2,7-dichlorodihydrofluorescein diacetate, DCF-DA). Scale bar 10 μm.

The H_2_O_2_ production was verified *in vivo* by staining with CM-H_2_DCFDA (DCF), a probe for intracellular H_2_O_2_ detection and simultaneously mark with the mitochondria-selective probe MitoTracker Red CMXRos (Molecular Probes) using confocal microscopy (Figure IIIB). It was found a green DCF signal that co-localized with MitoTracker Red in small (< 1μM diameter) circular-shaped structures, being the DCF signal more intense in the L-GaL-treated tissue (Figure IIIB). It corroborates that H_2_O_2_ was produced inside the mitochondria. Consistently, we found that the *in vivo* ROS staining was still detected in the presence of inhibitor of AOX, slightly decreased with Cytc oxidase’s inhibitor but almost fully disappeared in fruit tissue treated with antimycin A, rotenone and DPI (Figure IIIB). *In vivo* mitochondrial activity in fruit tissue was confirmed by observing the depletion of MitoTracker Red and DCF signals in presence of the mitochondrial uncoupler, CCCP (Data not shown).

We also demonstrate the lower production of H_2_O_2_ in fruit mitochondria from L-GalLDH-RNAi plant lines, which was consistent with an increased AOX respiration during AsA synthesis (supplementary data V). However, despite these mitochondria showed decreased L-GalLDH activity (lower Cytc reduction rate), their abilities to produce AsA and alter FAD redox status were similar to that of wild type fruit mitochondria (supplementary data V). It suggests that AOX pathway may sustain AsA synthesis in mitochondria with low L-GalLDH activity by reducing H_2_O_2_ level.

To further examine the role of alternative respiration during AsA synthesis, we performed an opposite experiment in which the AOX is previously activated before the treatment with L-GalL. To this, mitochondria were firstly treated with pyruvate, a known allosteric AOX activator, and subsequently L-GalL was added to inhibit alternative respiration. Surprisingly, the AOX respiration (7.6 nmol O_2_ mg min^−1^protein^−1^) was not inhibited in the presence of pyruvate (supplementary data VI). Moreover, this lack of inhibitory effect was not related to lower L-GalLDH activity given that its capacity to reduce Cytc remains in presence of pyruvate. However, the higher AOX capacity correlates with lower H_2_O_2_ production and enhanced AsA synthesis (supplementary data VI).

The possibility of that AsA synthesis leads to shifts in the overall functional status of cell was tested by performing a comparative proteomic analysis between untreated and L-GalL-treated papaya fruit tissue. Of the set of 53 proteins identified, 24 (45%) and 29 (55%) were up-regulated and down-regulated by L-GalL, respectively (Table II). Possible roles of these proteins will be discussed later with regards to an involvement of AsA synthesis in ROS and energy metabolism as well as in the plant responses to abiotic and biotic stresses.

**Table II.** Summary of proteins differentially regulated by L-GalL in papaya fruit mesocarp.

| Identification number | Blast2GO description | Fold change L-GalL/Control |
| --- | --- | --- |
| **Up-regulated proteins\*** |  |  |
| pacid=16406756 | 40S ribosomal S8 | 7.2 |
| pacid=16407487 | pyruvate kinase cytosolic-like isoform X1 | 7.1 |
| pacid=16427581 | probable thimet oligopeptidase isoform X3 | 3.5 |
| pacid=16431084 | SEC14 cytosolic factor family phosphoglyceride transfer family isoform 2 | 2.6 |
| pacid=16404296 | 40S ribosomal S2-2-like | 2.3 |
| pacid=16425273 | glycerophosphodiester phosphodiesterase chloroplastic-like | 2.3 |
| pacid=16421901 | Hydroxyproline-rich glyco family isoform 2 | 2.1 |
| pacid=16423413 | 60S ribosomal L8-3 | 2.1 |
| pacid=16408435 | plasma membrane ATPase 4 isoform X1 | 2.1 |
| pacid=16423505 | Phospholipase D delta | 2.1 |
| pacid=16415693 | clathrin light chain 1-like | 2.0 |
| pacid=16407929 | sirohydrochlorin chloroplastic-like | 2.0 |
| pacid=16409797 | transport SEC23-like | 1.9 |
| pacid=16409547 | probable thiol methyltransferase 2 isoform X3 | 1.9 |
| pacid=16404282 | 60S ribosomal L23A | 1.8 |
| pacid=16415376 | NADH dehydrogenase subunit 7 (mitochondrion) | 1.8 |
| pacid=16421330 | ferritin- chloroplastic-like | 1.7 |
| pacid=16410499 | argonaute 1 | 1.7 |
| pacid=16417105 | probable low-specificity L-threonine aldolase 1 | 1.6 |
| pacid=16422730 | cell division cycle 48 homolog | 1.6 |
| pacid=16426064 | probable glutathione S-transferase | 1.6 |
| pacid=16423032 | 40S ribosomal S3a | 1.6 |
| pacid=16405013 | GATA zinc finger domain-containing isoform 1 | 1.5 |
| pacid=16407528 | phosphoenolpyruvate carboxykinase [ATP] | 1.5 |
| **Down-regulated proteins** |  |  |
| pacid=16422946 | cysteine synthase | 0.7 |
| pacid=16429847 | L-ascorbate peroxidase cytosolic | 0.7 |
| pacid=16426651 | Succinate dehydrogenase 5 | 0.7 |
| pacid=16406290 | probable fructokinase- chloroplastic | 0.6 |
| pacid=16415704 | alpha-galactosidase family | 0.6 |
| pacid=16417606 | nascent polypeptide-associated complex subunit alpha 1 | 0.6 |
| pacid=16431407 | thioredoxin-dependent peroxidase | 0.6 |
| pacid=16427996 | probable polygalacturonase isoform X3 | 0.6 |
| pacid=16413137 | cysteine ase RD21A-like | 0.6 |
| pacid=16426375 | glyoxylate succinic semialdehyde reductase 1 | 0.6 |
| pacid=16413886 | nucleoside diphosphate kinase B | 0.6 |
| pacid=16406032 | leucine--tRNA cytoplasmic | 0.6 |
| pacid=16428645 | 14 kDa zinc-binding | 0.6 |
| pacid=16414991 | E3 ubiquitin- ligase RNF25 | 0.5 |
| pacid=16406747 | NAD(P)H dehydrogenase (quinone) FQR1 | 0.5 |
| pacid=16412645 | 26S protease regulatory subunit 6B homolog | 0.5 |
| pacid=16418146 | PREDICTED: uncharacterized protein LOC105784633 | 0.5 |
| pacid=16411412 | GTP-binding SAR1A | 0.5 |
| pacid=16420462 | heat shock 70 kDa | 0.5 |
| pacid=16413904 | GTP-binding YPTM2 | 0.4 |
| pacid=16414185 | 3-hydroxyisobutyryl- hydrolase mitochondrial isoform X1 | 0.4 |
| pacid=16422671 | MLP 43 | 0.4 |
| pacid=16405578 | alpha-1,4-glucan- synthase [UDP-forming] 2 | 0.4 |
| pacid=16425822 | cysteine ase inhibitor 5 | 0.4 |
| pacid=16424442 | cell division cycle 48 homolog | 0.2 |
| pacid=16426123 | trehalose-6-phosphate synthase | 0.1 |
| pacid=16430375 | dirigent 22-like | 0.06 |
| pacid=16425257 | 10 kDa chaperonin-like | 0.05 |

\* Up-regulated and down-regulated proteins represent those having a fold change of at least 1.5 (ANOVA, $p < 0.05$ ) relative to the treatment without L-GalL (control). As criteria for reproducibility, only proteins identified with differential abundance were included in the table. The bioinformatic analysis used the *Carica papaya* v. 0.4 protein databank from Phytozome and the functional annotation was done using Blast2Go software v.3.4.

### Regulation of seedling emergence by L-GalLDH is associated with altered ATP content

To get further insights about the physiological role of L-GalLDH on seedling establishment (an energy-demanding process), we evaluated AsA synthesis and ATP content in both L-GalLDH-RNAi lines and wild type in germinating seeds and in seedlings that reach the autotrophy capacity. Wild type and L-GalLDH-RNAi seedlings showed similar ability to synthesize AsA, but, wild type ones contain less ATP in dark (Figure 4A). Interestingly, wild type germinating seeds also had a little less ATP content (data not shown). Treatment of seeds with L-GalL inhibited wild type seedling emergence and consequently, they showed shorter seedlings one-week post germination (Figure 4B). However, this inhibitory effect of L-GalL was not evident in L-GalLDH-RNAi lines and these seedlings elongated faster (Figure 4B). Under the growth conditions used in the experiment, these differences in size between wild type and plant lines disappeared when seedlings became larger. In fact, at 30-days-old stage, wild type seedlings had higher size (Figure 4B). Nonetheless, carbon dioxide fixation and biomass were comparable between wild type and L-GalLDH-RNAi plant lines at 30 days after germination (Figure 4C).

**Figure 4A.**
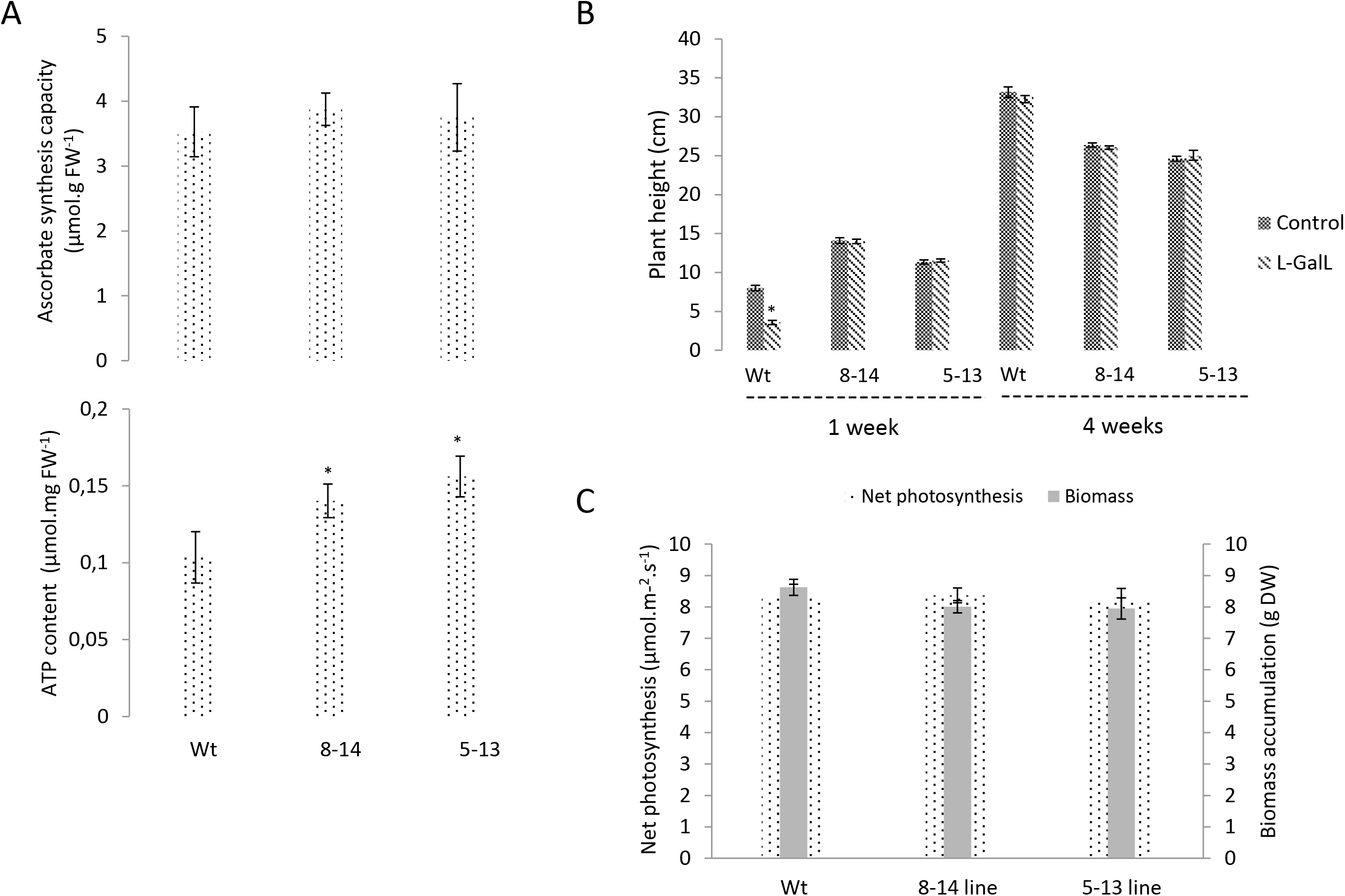
Ascorbate synthesis capacity and ATP content of leaf tissue (in dark) determined in leaf tissue from four weeks-days-old wild type, 8-14 and 5-13 transgenic lines. **4B**. Effect of seed treatment with 20mM L-GalL on seedling growth from wild type, 8-14 and 5-13 transgenic lines measured one-week and four-weeks following the chemical treatment. **4C**. Net photosynthesis of fully expanded leaves (Dotted line) and dry weight of aboveground tissue (Grey line) determined in wild type, 8-14 and 5-13 transgenic lines at the four-weeks growth stage. All bars are means ± standard error from three independent experiments.

## DISCUSSION

Over many years, it has been believed that plants synthesize AsA basically to produce this powerful antioxidant molecule, which has multiple functions (Smirnoff, 2018). The localization of the L-GalLDH enzyme within mitochondria has supported the obvious paradigm that AsA synthesis exists for producing AsA, which promotes ROS scavenging. We unexpectedly found that AsA synthesis triggers ROS content and down-regulates energy supply. Most specifically, the electron flux derived from L-GalLDH activity is poorly used for the generation of proton gradient and consequently ATP supply. Instead, AsA synthesis interferes with the conventional electron flux associated with the oxidation of reducing equivalents (NADH/FADH2). Surprinsigly, this inhibitory effect of the respiratory flux had an unknown characteristic in plants; it is the possibility of activation of a reverse electron flux during AsA synthesis. Therefore, a novel perspective about the bioenergetics of plant mitochondria mayarise: Plants might use the AsA synthesis not only to accumulate this antioxidant but also to induce a new manner to alter the mitochondrial respiratory activity and their redox balance.

Unexpectedly, we also see that the known mitochondrial AsA synthesis pathway, which transfers electrons through Cytc (Bartoli et al., 2000) can also transfer electrons towards ubiquinone. It was concluded that AsA synthesis pathway is branched and can function without the need of Cytc oxidase activity. Data show that FAD, which is an electron carrier for L-GalLDH (Leferink et al., 2009) might be enrolled in the electron transfer. Intriguingly, complex II also requires FAD as internal electron carrier for reducing ubiquinone (Schertl and Braun, 2014) and the proteomic analysis revealed a down-regulation of subunit 5 of complex II, which has unknown function so far in plants (Huang and Millar, 2013) but in yeast is required for FAD incorporation (Hao et al., 2009). These facts may suggest that AsA synthesis down-regulates the respiratory transport by interfering with the electron entry through complex II. The existence of a branched AOX-dependent pathway that would interact with the complex II during AsA synthesis may have practical implications. If electrons derived from AsA synthesis can flow via FAD towards the ubiquinone without passing through Cytc, new alternative assays for quantifying L-GalLDH activity might be developed.

Similarly to the demonstrated role of AsA synthesis, the AOX pathway can also decline ATP production; however, the later pathway does not reduce the mitochondrial capacity to utilize reducing equivalents (Schertl and Braun, 2014). This difference may be strongly linked with the close inter-relationship between AOX and L-GalLDH expressions (Bartoli et al., 2006). Likely, when plants need to decrease mitochondrial ATP generation, the L-GalLDH enzyme could be expressed to inhibit the respiratory electron flux unlike the AOX pathway, which maintains a significant part of such flux intact, contributing to energy loss. Therefore, the AsA synthesis may be activated to avoid the loss of energy when the need for mitochondrial ATP is low.

Our study also shows that the generation of hydrogen peroxide (H_2_O_2_) during AsA synthesis is crucial for determining AsA level. Paradoxically, AsA is needed for H_2_O_2_ elimination (Smirnoff, 2018) and L-GalLDH and AOX activities are sensitive to H_2_O_2_ (Leferink et al., 2009). Likely, H_2_O_2_ also plays a role in inactivating AOX pathway during AsA synthesis. AOX inactivation occurs by the formation of a disulfide bridge between the two cysteine residues in the AOX dimer under oxidizing conditions (Kühn et al., 2015) and it has been proposed as a regulatory mechanism for AOX activity (Selinski et al., 2018). AOX plays a role in minimizing ROS load in plants (Dahal and Vanlerberghe, 2017), which may explain why AOX activity showed to be necessary for AsA synthesis and L-GalLDH activity. An excessive hydrogen peroxide level during AsA synthesis might be prevented by coordinated AOX and L-GalLDH protein expressions. In line with this hypothesis, previous results showed a synergism between AOX expression and AsA accumulation in Arabidopsis plants under light (Bartoli et al., 2006). Accordingly, AOX pathway may function to protect plants from the pro-oxidant effect of AsA synthesis.

We also hypothesized that the H_2_O_2_ coming along with AsA synthesis may play a role in defining cell’s redox status. Mitochondria can exert a strong control over the redox balance of the cell (Noctor et al., 2007). As a signal molecule, the H_2_O_2_ produced during AsA synthesis would diffuse to the extra-mitochondrial environment, being a major regulator of redox signaling and protein expression. Our proteomic data revealed regulations of ROS-related enzymes beyond mitochondria during AsA synthesis. Notably, the cytosolic ascorbate peroxidase (cAPX), which utilizes AsA and hydrogen peroxide as substrates (Davletova et al., 2005) was highly responsive. Most of the identified proteins were previously involved in regulatory redox cascades related with hormonal signaling, defense/detoxification, protein folding and transcriptional/translational regulation, membrane/protein trafficking and degradation, programmed cell death (PCD), fruit ripening as well as in stress responses to high light, hypoxia, drought, iron storage, sulfur metabolism and phosphorus starvation (Westlake et al., 2015). The whole picture is consistent with the hypothesis of that these changes of cellular redox state are involved in a retrograde signal transduction associated with mitochondrial AsA synthesis.

In addition, retrograde signal associated with AsA synthesis may be regulating sugar and lipid catabolism and cytosolic ATP provision. We note regulations of enzymes linked with extra-mitochondrial ATP generation through glycolysis and sugar metabolism in cytosol and/or cell wall. Pyruvate kinase, phosphoenolpyruvate carboxykinase, threonine aldolase, fructokinase, α-galactosidase, trehalose-6-phosphate synthase and polygalacturonase, which were identified in the proteomic, have been implicated (Schluepmann et al., 2003; Umbach et al., 2006). Consistently, AsA-related mutants present altered sugar metabolism (Alhagdow et al., 2007) and our results showed effects on initial growth and ATP content. AsA has been linked with cell growth regulation in plants (Arrigoni et al., 1997). Studies with low ascorbate Arabidopsis mutants (*vtc-1* and *vtc2-1*) revealed that these plants had limited growth (Plumb et al., 2018). Thus, aplausible hypothesis may be that the rate of AsA synthesis controls the supply of mitochondrial energy for growth.

Based on these unanticipated findings, we believe that the wide variability in the synthesis rate of this powerful anti-oxidant molecule in different plant sources, species, during lifecycle and environmental situations might reflect a distinct manner of regulation of plant capacity for adapting their mitochondria to fluctuating energy demands and redox status.

## MATERIALS AND METHODS

### Plant material

Cherry tomato plants from wild type genotype *(Solanum lycopersicum* ‘West Virginia 106’) and the tomato *P_35S_:Slgalldh^RNAi^* silenced lines 5 and 8 (Alhagdow et al., 2007) were all grown under standard greenhouse conditions. Tomato fruits were harvested at green-mature stage. For papaya and strawberry, green-mature fruit (*Carica papaya*, ‘Golden’ cultivar) and red-mature fruit (*Fragaria vesca*, ‘Oso Grandi’ cultivar) were obtained from local suppliers. When fully-ripe fruits were needed, post-harvest ripening were performed in chambers with controlled temperature (25°C ± 1°C) and relative humidity (85% ± 5%) during nine days. The degree of fruit ripening was assessed by the changes in skin color and pulp firmness (Oliveira et al., 2015).

### Mitochondria isolation and purification

Leaf mitochondria were purified following a Percoll density-gradient method as previously described (Keech et al., 2005). Fruit mitochondria were isolated using fruit mesocarp of papaya, tomato and strawberry with defined ripening characteristics, and isolations were performed following general procedures, modified for fruit mitochondria, which were described in (Oliveira et al., 2017).

### Respiratory measurements of isolated mitochondria

Immediately after purification, respiration measurement was assessed by O_2_ exchange in a Clark electrode (Oxytherm system, Hansatech, UK) using fresh and intact mitochondria (around 20-80 μg mitochondrial protein) resuspended in reaction medium (0.35 M mannitol,10 mM MOPS, 10 mM KPO4, 10 mM KCl, 5 mM MgCl2, and 0.5% (w/v) defatted BSA at pH 7.2 and 25°C). The AOX capacity was assessed by measuring the O_2_ consumption rate sensitive to n-propyl gallate (inhibitor of AOX pathway) in the presence of 3 mM of KCN or NaN3 (Oliveira et al., 2015). The respiratory control rate (RCR) and ADP:O ratio of intact mitochondria were determined using 10 mM malate and respiration was calculated as the O_2_ uptake rate of the coupled mitochondria in non-phosphorylating state 4, i.e., after consumption of all ADP added in the absence of any inhibitors, as described in previous works (Oliveira et al., 2015). When indicated, mitochondria were energized with distinct respiratory substrates (5 mM L-GalL, 8 mM NADH, 10 mM malate, 20 mM glutamate, 5 mM AsA) and oxygen uptake was recorded in presence of AOX and COX inhibitors, salicylhydroxamic acid (SHAM) and NaN3, respectively. When needed, mitochondria were also treated with 60 μM of CCCP (carbonyl cyanide *m*-chlorophenyl hydrazone) used as mitochondrial uncoupler of proton gradient (Oliveira et al., 2015).

### Measurement of H_2_O_2_ production by purified mitochondria

The rate of H_2_O_2_ formation was determined using Amplex Red/horseradish peroxidase (HRP) assay, in a 96-well microplate using a Chameleon Microplate reader (HIDEX), through the detection of the highly fluorescent resorufin, as described previously (Gleason et al., 2011). The concentrations of Amplex Red and HRP in the incubation medium (100 μL final volume per well) were 50 μM and 0.1 U/mL, respectively. The mixture also contained about 10-50 μg of mitochondrial protein prepared in 10 mM MOPS, 10 mM KCl, 5 mM MgCl2 at pH 7.2 and 25°C. For testing respiratory inhibitors/activators, the reaction was supplemented with the defined chemical compounds before adding L-GalL. The chemicals included rotenone (complex I inhibitor), salicylhydroxamic acid, SHAM (AOX inhibitor), pyruvate (AOX activator), antimycin A (complex III inhibitor), diphenylene iodonium, DPI (a flavin-containing oxidase inhibitor) and NaN_3_ (complex IV inhibitor). When SHAM was tested, the incubation medium had a higher HRP concentration (0.6 U/mL). The reaction was initiated by adding defined L-GalL concentrations into incubation medium and the L-GalL-dependent fluorescence change was recorded with 570 nm excitation and 585 nm emission wavelengths during 15 min. Fluorescence backgrounds of control reactions containing the tested chemicals without L-GalL were allowed to stabilize for two minutes before L-GalL was added to start reaction. To calculate the H_2_O_2_ level, these backgrounds were subtracted from all fluorescence measured after adding L-GalL. Importantly, as the different compounds used affect the basal fluorescence signal, calibration of all background reactions was done using the same compound concentrations and mitochondrial preparations. The H_2_O_2_ production was expressed as the corrected change in fluorescence in relation to the corresponding controls.

### In vivo detection of ROS formation

Mesocarp discs from green-mature papaya fruit were treated with 5 mM L-GalL in 50 mM MOPS buffer pH 7.0 with or without defined respiratory inhibitors that included 2 mM NaN_3_, 1 mM SHAM, 20 μM rotenone, 2 μM antimycin A and 5 mM DPI. The treatment of fruit mesocarp with chemicals lasted two hours. Control tissue was subjected to the same procedure without the addition of L-GalL. Afterwards, tissues were incubated in dark for 10 min at 25°C with 500 nM of carboxy-H_2_DCFDA (Molecular Probes) and 50 nM of MitoTracker Red CMXRos (Invitrogen). Carboxy-H_2_DCFDA permeates membranes and is retained by cells after cleavage of the acetate moiety by cellular esterases and fluorescence develops upon oxidation of the dye by ROS. MitoTracker Red is a membrane potential-sensitive probe that accumulates into mitochondria (Fricker and Meyer, 2001). Stock solutions of the dye were prepared in dimethyl sulfoxide and kept in the dark at −80°C. Before microscopy, samples were quickly rinsed in 50 mM MOPS buffer pH 7.0 to remove any excess of dyes that had not penetrated into the tissue. Microscopy observations were performed using a confocal laser scanning microscope system (Leica TCS SPE laser scanning confocal microscope) using a 63x with 1.4 numerical aperture objective. Carboxy-H_2_DCFDA signal was visualized with excitation at 492 nm and emission at 527 nm. MitoTracker Red signal was visualized with excitation at 579 nm and emission at 599 nm. All images were obtained digitally quickly after excitation and all sections observed under the same microscopy parameters by z series stacking. The experiment was repeated twice and led to similar results.

### Determination of AsA synthesis capacity in isolated mitochondria and plant tissue

Total AsA (reduced and oxidized AsA) was determined as previously described in (Mazorra et al., 2014), with modifications. Briefly, fresh purified mitochondria (10-40 μg) were incubated in 5 mM of L-GalL dissolved in 50 mM TRIS buffer pH 7.8 (in absence of mannitol) for 15, 30, 60 and 120 min. When indicated, incubations with L-GalL were performed in presence of respiratory protein inhibitors (rotenone, antimycin A, and DPI) or the AOX activator (pyruvate). For each treatment, corresponding control samples without L-GalL were also included. Reactions were stopped with 5% (v/v) trifluoroacetic acid (TFA), centrifuged at 10.000g at 4°C for 10 min, pellet discarded and supernatant neutralized with drops of 100 mM K_2_HPO_4_ pH 10. Then, samples were reduced with 5 mM dithiothreitol (DTT) for 5 min, quickly filtrated and finally 20 μL were injected and separated by HPLC (Shimadzu, Kyoto, Japan) using a Spherisorb ODS C-18 column equilibrated with 100 mM phosphate buffer pH 3.0. Runs were performed at 1 mL min^−1^ flux rate at 25 °C. AsA peak was detected at 254 nm with a coupled UV-detector. The AsA synthesis capacity of mitochondria was expressed as the total AsA produced per mitochondrial protein (mg) per time (min). Determination of AsA content in plant tissues was performed as previously described (Bartoli et al., 2006). The AsA synthesis capacity was expressed as total AsA per fresh weight (mg).

### Measurement of cytochrome c reduction capacity

The L-GalL-induced Cytc reduction capacity was assessed following procedure described in (Ôba, et al., 1995) with modifications. Purified mitochondria (40-60 μg) were assayed in medium containing 50 mM TRIS buffer pH 7.8, 5% (v/v) Triton X-100, 5 mM Cytc and the reaction was started by adding 5 mM of L-GalL. The formation of reduced cytochrome c was spectrophotometrically monitored at 550 nm every 10 seconds for two minutes. The extinction coefficient of Cytc (21.1 mM^−1^ cm^−1^) was used for calculating the reaction rate. As indicated, different respiratory inhibitors alone or combined were added into reaction mixture before the start of reaction. The capacity to reduce Cytc was expressed as the amount of Cytc reduced per mitochondria protein per min.

### Assessment of changes in FAD redox state

Changes in FAD redox state can be detected based on the fact of FADH_2_ and FAD are a redox pair but only oxidized FAD is fluorescent and can be monitored without exogenous labeling at 450 nm excitation and 520 nm emission wavelengths (Kozioł, 1971). To this, purified mitochondria (20-40 μg) were resuspended in 0.6 mM FAD and 50 mM TRIS buffer pH 7.8 without mannitol. The basal fluorescence was allowed to stabilize during 20 min and then 5 mM L-GalL was added to induce AsA synthesis. The change in fluorescence was recorded every 30 min in both reactions with or without L-GalL. To determine the effects of inhibitors on FAD fluorescence, when indicated, inhibitors were added and the fluorescence backgrounds were recorded. Then, changes in inhibitor-dependent FAD fluorescence in presence of L-GalL were determined. Alterations in FAD redox state were expressed as the difference between fluorescence (final minus initial) values within 30 minute intervals.

### Measurement of ubiquinol production capacity

To assess the ability of L-GalL to convert ubiquinone into ubiquinol, purified mitochondria (40-60 μg) were incubated with 5 mM of L-GalL, 15 μM of ubiquinone (UQ10) in 1 mL of 50 mM TRIS buffer pH 7.8 for 4 h in dark. Incubations were done in the presence of 3 mM NaN_3_, 1 mM SHAM to avoid ubiquinol re-oxidation. Controls without L-GalL were also included. Extractions were performed based on procedure published by (Wagner and Wagner, 1995) with modifications. Briefly, reactions were stopped with 10 % (v/v) TFA, extracted in shade with 600 μL of cold-hexane, vortexed for 1 min, centrifuged at 6.000g at 4°C for 5 min. Then, the upper hexane phase was collected, evaporated to dryness in a rapidVap evaporation system and finally the extracted UQH_2_/UQ was re-suspended in 500 μL of acetone. Samples were filtrated to analyze in HPLC. 20 μL were injected in reverse-phase C-18 column equilibrated with acetonitrile:etanol 3:1 (v/v). Runs were done at 1.5 mL min^−1^ flux rate and 40°C. The ubiquinol (UQH_2_10) peak was detected with a fluorescence detector at 290 nm (exc), 370 nm (em), as described in Yoshida et al. (2010). The ubiquinone reduction capacity was expressed as the ubiquinol/ubiquinone ratio determined after incubation in the presence of L-GalL over 4 hours.

### Assessment of NADH production capacity by isolated mitochondria

The NADH production was based on the difference in the absorption spectra of NAD^+^ and NADH. The NADH shows absorption maxima at 340 nm but NAD^+^ does not absorb light at 340 nm (Renault et al., 1982). In brief, mitochondria were osmotically-broken by incubating into 10 mM MOPS (without mannitol) pH 7.2, 0.1 mM EDTA during 10 min. Then, a reaction mixture was prepared consisting of 5 mM L-GalL, 20 mM NAD^+^, 3 mM azyde and 1 mM SHAM, allowed to react during 30 min at 25°C and NADH absorbance (extinction coefficient NADH = 6.220 M^−1^ cm^−1^) was read with UV/Vis spectrophotometer. Control mixture had the same components, except L-GalL. The relative NADH production was determined as the difference in absorbance at 340 nm following incubation for 30 min.

### Measurement of H^+^-ATPase pumping activity

The H^+^-ATPase activity was measured using sub-mitochondrial particles obtained from intact mitochondria treated or not with L-GalL. Particles were prepared by sonication, as described in (Ragan et al., 1987), with some modifications. Briefly, intact mitochondria (200 μg) were incubated in 10 mM MOPS containing 0.35 M mannitol with or without 5 mM L-GalL over 60 min at 25°C. Following treatment, mitochondria were sonicated by 6-10 s pulses with 30 s intervals and supernatant ultra-centrifuged and the resulting pellet (sub-mitochondrial particles) were re-suspended into the same buffer. Then, sonicated-disrupted mitochondria solution was incubated with 1 mM ATP and its capacity to hydrolyze ATP was monitored by measuring the release of inorganic phosphate (P_i_) colorimetrically at 720 nm, as previously reported in (Subbarow, 1925). H^+^-ATPase activity was expressed as the amount of released Pi during 1 min into the reaction medium.

### Measurement of ATP level

To determine the mitochondrial capacity to synthesize ATP during ascorbate biosynthesis, 50 μg of freshly purified intact mitochondria were incubated in medium containing 10 mM MOPS, 150 mM sucrose, 7.5 mM KCl, 5 mM MgCl_2_, 7.5 mM KPi and 5 mM of L-GalL during 10 min. Then, reaction was initiated adding 10 mM malate and 50 μM of ADP and allowed to incubate at 25°C during 30 min. Control reactions were run in absence of ADP and malate. Mitochondrial ATP was extracted by boiling the samples for 15 min. After centrifugation at 9000g for 15 min, ATP content in 200 μL of supernatants was determined by the bioluminescent assay based on luciferin-luciferase method (Sigma-Aldrich, FLAA) using a luminescence spectrophotometer (RF5301PC, Shimadzu) at 560 nm of emission wavelength. The reaction was initiated by addition of 20 μL of ATP assay mix (Sigma-Aldrich, FLAAM) in a mixture containing 1800 μL of ATP assay buffer (10 mM MgSO_4_, 1 mM DTT, 1 mM EDTA, 100 μg.mL^−1^ bovine serum albumin, and 50 mM tricine buffer salts, pH 7.8) and 200 μL of sample. The reaction was monitored for 1 min. Calibration curve was performed previously in the same conditions with ATP standard solutions ranging from 0.1 to 10 μmol.

To determine ATP content in plants, 600-900 mg of aboveground plant tissue were collected at night-time (two hours before lighting) and were quickly incubated at 100°C for 15 min in 1 mL of boiled water, as described in (Yamamoto et al., 2002). Tissues were homogenized at 4°C and then centrifuged at 9000g for 15 min at 4°C. Supernatant (200 μL) was used for ATP quantification, as described above.

### Western blot analysis

Mitochondrial proteins were reduced using 2.5% (v/v) 2-mercaptoethanol into sample buffer, loaded onto one-dimensional SDS/PAGE gels and run following standard procedures. For the detection of oxidized AOX, mitochondrial proteins were prepared in absence of 2-mercaptoethanol. Molecular weight markers (24-102 kDa, GE Healthcare) were used and equal loading of gels (25 μg protein) was checked by Ponceau staining. The proteins were transferred to a nitrocellulose membrane (Hybond ECL, Amersham/GE Healthcare); the membrane was blocked in nonfat milk 5% overnight at 4°C and antibodies against L-GalLDH and AOX (commercially provided by Agrisera) were used in dilutions 1:500. Then, the membranes were washed three times in PBS buffer with milk 5%, incubated with goat anti-rabbit secondary antibody for 2h, and subsequently washed in PBS buffer three times. Results were visualized by chemiluminiscence with a ECL Western Blotting Detection System (Amersham/GE Healthcare) and quantified using ImageJ densitometric software (https://imagej.nih.gov/ij/).

### Experimental design for comparative proteomic analysis

Papaya fruit mesorcarp discs (500 mg fresh weight) were treated with 5 mM of L-GalL, 50 mM MOPS buffer pH 7.0 for two hours at 25°C. A control sample without L-GalL was also incubated under the same conditions. Then, three independent samples of proteins were extracted from each treatment. Procedures for protein extraction, digestion and mass spectrometry analysis are performed following previous works described in (Heringer et al., 2017).

### Bioinformatic analysis

Progenesis QI for Proteomics Software V.2.0 (Nonlinear Dynamics, Newcastle, UK) were used to spectra processing and database searching conditions. The analysis were performed using following parameters: Apex3D of 150 counts for low energy threshold, 50 counts for elevated energy threshold, and 750 counts for intensity threshold; one missed cleavage, minimum fragment ion per peptide equal to two, minimum fragment ion per protein equal to five, minimum peptide per protein equal to two, fixed modifications of carbamidomethyl (C) and variable modifications of oxidation (M) and phosphoryl (STY), and a default false discovery rate (FDR) value at a 4% maximum, peptide score greater than four, and maximum mass errors of 10 ppm. The analysis used the *Carica papaya* v. 0.4 protein databank from Phytozome (https://phytozome.jgi.doe.gov/). Label-free relative quantitative analyses were performed based on the ratio of protein ion counts among contrasting samples. After data processing and to ensure the quality of results, only proteins present in 3 of 3 runs were accepted. Furthermore, differentially abundant proteins were selected based on a fold change of at least 1.5 and ANOVA (*P* ≤0.05). Functional annotation was performed using Blast2Go software v. 3.4 (Conesa et al., 2005).

### Seed treatment, plant growth and photosynthesis

To determine growth of tomato plants with induced AsA synthesis, seeds of wild type and L-GalLDH-RNAi plant lines were subjected to imbibition treatment with 20 mM GalL for 6 hours. In parallel, control seeds were treated in absence of L-GalL. Imbibited seeds were sown on soil pots filled with commercial substrate and irrigated with Hoagland solution. Then, seedlings (one per pot) were grown for four weeks at a growth chamber at 25°C with a 16-h photoperiod at a light intensity of 500 μmol m^2^ s^−1^. Height and biomass of plants were determined at given time points. Dry weights were determined by drying aboveground tissue in an air circulation oven at 80°C for one week. Height was determined by measuring the distance from the ground to the top of canopy. At four-weeks after sowing, instantaneous gas exchange measurements were done on six recently fully expanded leaves in the upper part of the wild type plant and L-GalLDH-RNAi plant lines. Measurements were taken between 2-4 hours after the start of light period using a gas exchange system (LiCOR, Biosciences, Lincoln, NE, USA). Determinations of CO_2_ assimilation were performed at light intensity 500 μmolm^−2^s^−1^, 400 ppm CO_2_ and temperature 24-26°C.

### Statistical analysis

Data from at least three independent biochemical experiments were averaged and subjected to ANOVA and, when needed, means were analyzed following Tukey test at *P* ≤ 0.05.

## ACKNOWLEDGEMENTS

The authors (V.S., M.C., C.F.B., A.R.F., M.G.S., and J.G.O.) wish to acknowledge the Conselho Nacional de Desenvolvimento Científico e Tecnológico (CNPq-Brazil) for research fellowships. The authors L.M.M.M. and A.J.D.C. were granted with a CAPES (Coordenação de Aperfeiçoamento de Pessoal de Nível Superior)-PNPD; G.M.C.S. was granted with a FAPERJ (Fundação Carlos Chagas Filho de Amparo à Pesquisa do Estado do Rio de Janeiro)-PD; D.B.S., A.S.H., L.A.S.P., and A.V.O. were granted with a CAPES-DSc; S.F.P. was granted with a CNPq-DSc, and T.R.O. and R.S.R. were granted with a FAPERJ-DSc. fellowship.

